# Managing Uncertainty in Metabolic Network Structure and Improving Predictions Using EnsembleFBA

**DOI:** 10.1101/077636

**Authors:** Matthew B. Biggs, Jason A. Papin

**Affiliations:** Department of Biomedical Engineering, University of Virginia, Charlottesville, VA, USA

## Abstract

Genome-scale metabolic network reconstructions (GENREs) are repositories of knowledge about the metabolic processes that occur in an organism. GENREs have been used to discover and interpret metabolic functions, and to engineer novel network structures. A major barrier preventing more widespread use of GENREs, particularly to study non-model organisms, is the extensive time required to produce a high-quality GENRE. Many automated approaches have been developed which reduce this time requirement, but automatically-reconstructed draft GENREs still require curation before useful predictions can be made. We present a novel ensemble approach to the analysis of GENREs which improves the predictive capabilities of draft GENREs and is compatible with many automated reconstruction approaches. We refer to this new approach as Ensemble Flux Balance Analysis (EnsembleFBA). We validate EnsembleFBA by predicting growth and gene essentiality in the model organism *Pseudomonas aeruginosa* UCBPP-PA14. We demonstrate how EnsembleFBA can be included in a systems biology workflow by predicting essential genes in six *Streptococcus* species and mapping the essential genes to small molecule ligands from DrugBank. We found that some metabolic subsystems contribute disproportionately to the set of predicted essential reactions in a way that is unique to each *Streptococcus* species. These species-specific network structures lead to species-specific outcomes from small molecule interactions. Through these analyses of *P. aeruginosa* and six *Streptococci*, we show that ensembles increase the quality of predictions without drastically increasing reconstruction time, thus making GENRE approaches more practical for applications which require predictions for many non-model organisms. All of our functions and accompanying example code are available in an open online repository.

**Author Summary:** Metabolism is the driving force behind all biological activity. Genome-scale metabolic network reconstructions (GENREs) are representations of metabolic systems that can be analyzed mathematically to make predictions about how a biochemical system will behave as well as to design biochemical systems with new properties. GENREs have traditionally been reconstructed manually, which can require extensive time and effort. Recent software solutions automate the process (drastically reducing the required effort) but the resulting GENREs are of lower quality and produce less reliable predictions than the manually-curated versions. We present a novel method (“EnsembleFBA”) which overcomes uncertainties involved in automated reconstruction by pooling many different draft GENREs together into an ensemble. We tested EnsembleFBA by predicting the growth and essential genes of the common pathogen *Pseudomonas aeruginosa*. We found that when predicting growth or essential genes, ensembles of GENREs achieved much better precision or captured many more essential genes than any of the individual GENREs within the ensemble. By improving the predictions that can be made with automatically-generated GENREs, we open the door to studying systems which would otherwise be infeasible.

## Introduction

Metabolism is the driving force behind the wondrous flurry of biological activity carpeting our planet. An organism’s metabolism is determined by the metabolic enzymes encoded in its genome, the chemical reactions catalyzed by those enzymes, and whether or not those enzymes are actively expressed [1]. The simplest bacteria have hundreds of metabolic enzymes, while the most complex eukaryotes have thousands. The products of these enzymatic reactions serve as substrates for other reactions, such that the chemical transformations carried out in a cell can be represented as a vast network [2]. Mass and energy flow through such networks, transforming environmental inputs into the building blocks of life. Every species has a unique metabolic network driving its growth and interaction with the environment.

Genome-scale metabolic network reconstructions (GENREs) are formal representations of metabolic networks [3]. GENREs serve as a comprehensive collection of metabolic knowledge about a particular organism and they are amenable to mathematical analysis [4]. The process of reconstructing a GENRE takes months to years, but the reconstruction process often leads to new discoveries [5]. Mathematical analysis of GENREs gives insight into how particular metabolic pathways are used by an organism, what substrates it can utilize, which of its genes are essential in a given environment, how a metabolic network can be engineered to produce more of a desired product, or which enzymes within the network should be targeted in order to halt growth in an organism [6–8]. The reconstruction and analysis of GENREs for single species has greatly contributed to our understanding of microbes and our ability to engineer them. Recently, analyses have been developed which predict metabolic interactions between microbes [9,10]. However, the application of these recent analyses has been greatly limited by the large investment in time required to reconstruct a useful GENRE. Many microbial communities of interest consist of hundreds of species [11,12]. It is decidedly impractical to spend decades manually curating hundreds of GENREs.

Many automated methods have been developed for rapidly reconstructing more accurate GENREs [13– 16]. We present a novel ensemble method that is complimentary to these existing automated methods which we refer to as Ensemble Flux Balance Analysis (EnsembleFBA). EnsembleFBA pools predictions from many draft GENREs in order to more reliably predict properties that arise from metabolic network structure, such as nutrient utilization and gene essentiality (Fig 1). The primary benefits of this new method are that it relies on automatically-generated GENREs (which can be generated in a matter of minutes to hours) and yet produces more reliable predictions than individual GENREs within the ensemble. We implement and discuss one possible way of generating useful ensembles, but emphasize that other automated methods could be modified to generate useful ensembles.

**Fig 1.**
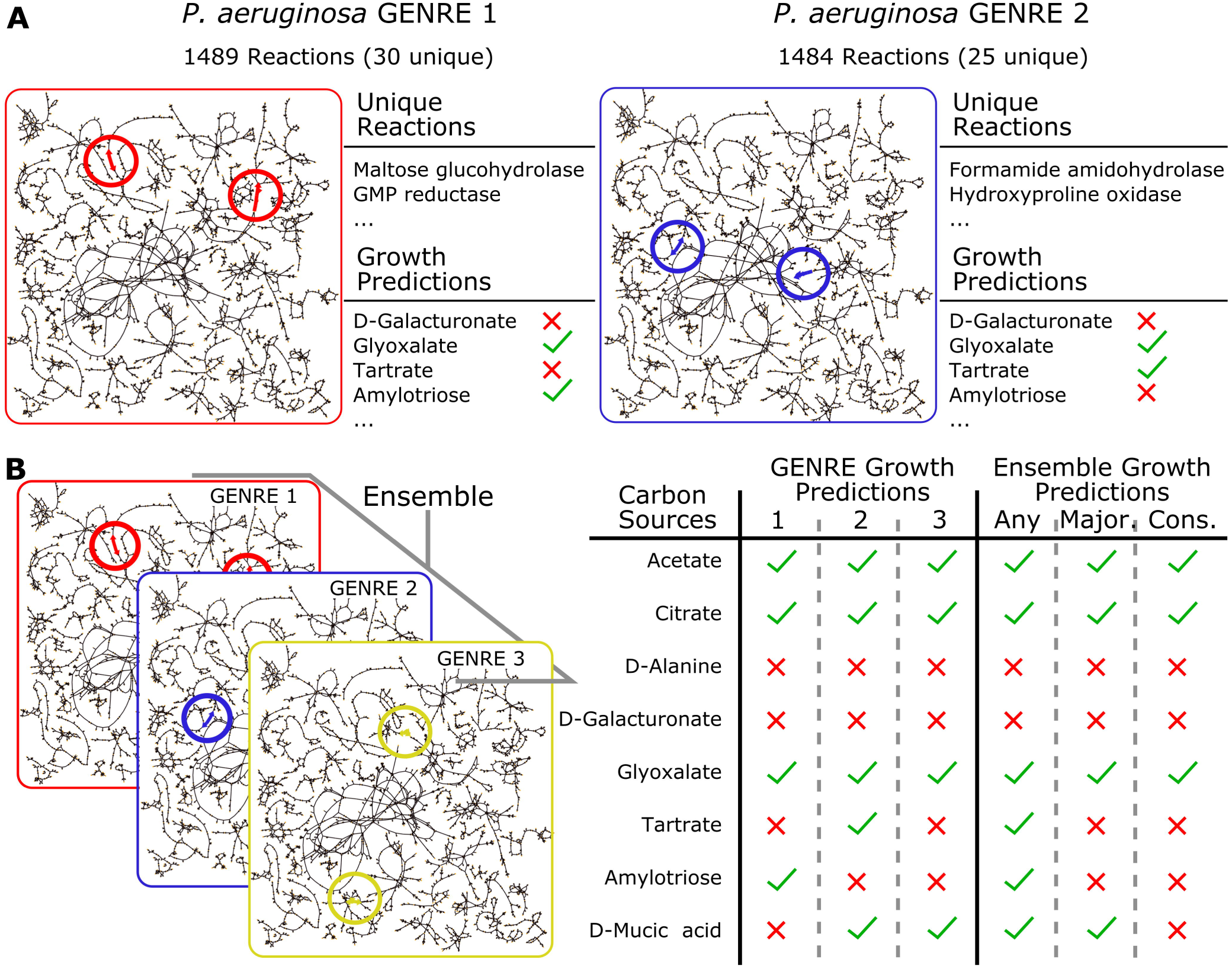
Alternative network structures can be analyzed collectively as an ensemble. **A**. Gap filling a network using the same media conditions, but in different orders, can lead to different network structures. Here we display two networks for *P. aeruginosa* gap filled with two permutations of the same 10 minimal media conditions. We highlight two unique reactions in each, and growth predictions which differ between the two. **B**. An ensemble can be created by collecting many alternative network structures which are all consistent with available data. Ensemble-level predictions are generated by treating the individual network predictions like votes. We used three qualitatively different decision thresholds: the “any” threshold requires that a single network predict growth; the “majority” threshold requires that a strict majority predict growth; the “consensus” threshold requires all networks within the ensemble to be in agreement. Note that the top five growth conditions result in the same prediction regardless of threshold, while the bottom three conditions result in threshold-dependent outcomes.

We begin by discussing a common GENRE curation procedure known as gap filling. We demonstrate that a global gap filling procedure does not perform any better than a sequential one. Instead, we introduce an ensemble approach to pool the many possible network structures resulting from different sequences of the input media conditions (Fig 1). We demonstrate that an ensemble reliably outperforms most of its constituent GENREs in terms of predicting growth and gene essentiality. By tuning the stringency of the voting threshold (e.g. requiring a majority of GENREs to agree vs. complete consensus) it is possible to achieve greater precision or recall than any of the constituent GENREs. We show how additional steps to increase the diversity among GENREs within the ensemble (e.g. reconstructing each member GENRE using subsets of the available data) can further improve recall. Furthermore, we found that incorporating negative growth information into our GENREs improved overall accuracy of the ensemble. We present proof of concept of the use of ensembles by predicting carbon source utilization and gene essentiality in *Pseudomonas aeruginosa*, a well-studied, clinically-relevant pathogen. We provide an example workflow using EnsembleFBA by predicting gene essentiality in six *Streptococcus* species and mapping the predicted essential genes to small molecules ligands in DrugBank. All of our data and code are available in an online repository, including example scripts to make adoption of EnsembleFBA easy. Our ability to make mechanistic predictions about complex cellular communities requires advances in the way we leverage the data available to us, and the way we handle uncertainty. Ensemble FBA is a novel tool that maintains the speed of automated reconstruction methods while improving predictions by intentionally managing uncertainty in network structures.

## Results

### Gap Filling Against Multiple Media Conditions in Different Orders Produces Different Network Structures

Gap filling is the process of identifying mismatches between computational predictions and experimental results, and identifying changes to the network structure which will bring the computational predictions into agreement with the experimental data. “Gaps” are missing reactions and can be filled by drawing from a database of possible metabolic reactions. Given that there are usually many mismatches between computational and experimental results, we demonstrate that simply changing the order in which computational results are brought into agreement with experimental can result in different network structures. For example, suppose that it is experimentally determined that a microbe can grow on glucose minimal media and sucrose minimal media, but the computational predictions do not match. Gap filling the GENRE against a representation of glucose minimal media first and sucrose minimal media second, may result in a different network in the end than if sucrose minimal media were first. In practice, the order of gap filling is arbitrary.

We implemented a custom gap fill algorithm based on the algorithms FASTGAPFILL and FastGapFilling (see Materials and Methods) [17,18]. We used the Model SEED biochemistry database as our “universal” reaction database from which to draw reactions for gap filling [13]. We used the Model SEED web interface to automatically generate a draft GENRE for *Pseudomonas aeruginosa* UCBPP-PA14 (without using the Model SEED gap filling feature). We gap filled this draft GENRE using 2, 5, 10, 15, 20, 25 and 30 media conditions that experimentally support growth, with 30 replicates for each. For example, we selected five media conditions at random and selected two random permutations of these conditions (in this case, there are 120 possible permutations). We gap filled in the order prescribed by the first permutation and then in the order of the second, and compared the resulting networks. We repeated the process 30 times, each time drawing a new set of five random media conditions and gap filling using two random permutations of those five media conditions. We found that even with as few as two media conditions, gap filling in a different order resulted in an average of 25 unique reactions per GENRE (Fig 2). As the number of media conditions increased, so did the average difference between the resulting GENRES.

**Fig 2.**
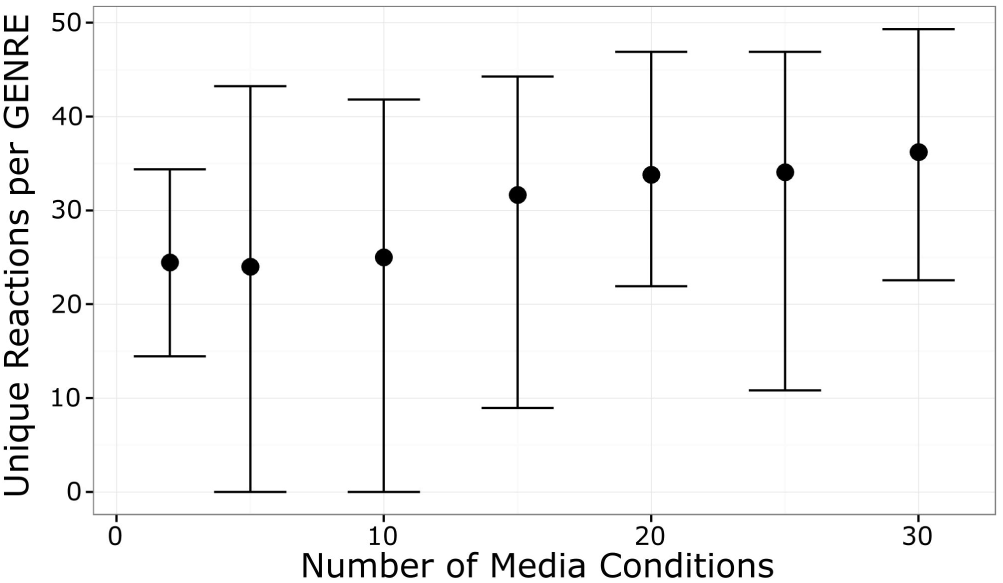
Gap filling in different orders leads to different network structures. Each error bar indicates an empirical 95% confidence interval from 30 simulations. For a single simulation, a set of media conditions was randomly selected (we simulated sets sizes of 2–30) and we gap filled a GENRE twice, using the same media conditions but in different orders. We compared the resulting pair of GENREs and we found that on average, GENREs within a pair contained an average of 25–35 unique reactions. The average number of unique reactions increased with the number of media conditions used to gap fill.

### “Global” Gap Filling Provides No Advantages Over a Sequential Approach

We hypothesized that rather than gap filling sequentially, perhaps a “global” gap fill approach would result in more parsimonious, biologically-relevant solutions without the ambiguity associated with changing the gap fill order. We extended our custom gap fill algorithm to identify a minimal set of reactions which could be added to a GENRE to permit growth in multiple media conditions simultaneously (see Materials and Methods). We started with the *P. aeruginosa* UCBPP-PA14 draft GENRE from the Model SEED and repeated the 30 replicates from 2 to 30 media conditions as above, but using the global gap fill approach that we developed (see Materials and Methods; Fig 3). We found that this global approach did not identify solutions that were any more parsimonious (Fig 3A), and lead to dramatic increases in solve times with increasing media conditions (Fig 2B). In order to determine whether the global solution was any more “biologically-relevant”, we also compared the ability of the global and sequential approaches to reconstruct a well-curated GENRE for *P. aeruginosa* UCBPP-PA14 called iPAU1129 (Bartell et al. In review). For each iteration (30 total), we removed 20% of reactions from iPAU1129 and used the sequential and global approaches to gap fill from the universal database using a random selection of five media conditions. The resulting networks were compared to the original iPAU1129, under the assumption that the most biologically-relevant approach would most faithfully reconstructed the curated GENRE, iPAU1129. We found no statistically significant difference between the two approaches (Fig 3C) (p-value = 0.63 by two-sided, paired Wilcoxon signed rank test).

**Fig 3.**
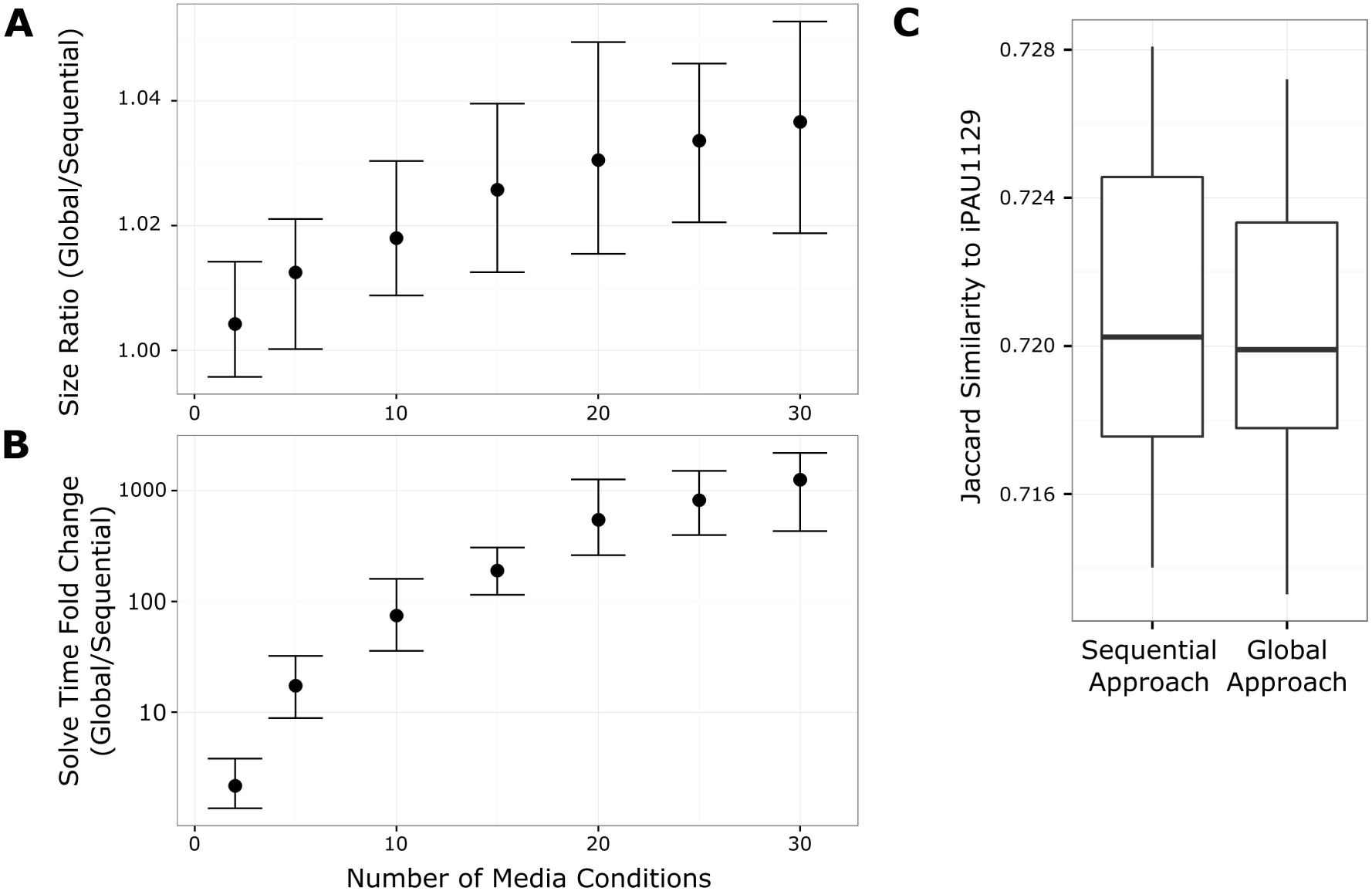
Results of global gap filling approach are no more parsimonious or biologically relevant. For each number of media conditions, we reconstructed 30 pairs of GENREs. For each pair, a set of media conditions was randomly selected and one GENRE was gap filled sequentially while the other was gap filled using a global approach. **A**. We found that the GENREs resulting from the global approach were slightly larger than those gap filled using the sequential approach, with an increasing size disparity as the number of media conditions increased. **B**. The global gap fill approach required significantly more time to run (note the log scale on the y-axis). The solve time increased quadratically with the number of media conditions, such that with 30 media conditions the average solve time for the global approach was ~1000 times greater than the sequential approach. Error bars in panels A and B represent empirical 95% confidence intervals. **C**. We compared the ability of the sequential and global approaches to replace reactions removed from a manually-curated GENRE for *P. aeruginosa* UCBPP-PA14, iPAU1129. For each replicate, we removed 20% of the reactions from iPAU1129 and applied the sequential and global gap filling approaches with the same set of randomly selected media conditions. We compared the reaction content of the gap filled GENREs with iPAU1129 using the Jaccard similarity metric. We found that there was no difference between the sequential and global approaches in terms of recovering the removed reactions (p-value = 0.63 by two-sided, paired Wilcoxon signed rank test). Box plots indicate quartiles of the distributions.

### Collecting Many Alternative Network Structures into an Ensemble Results in Improved Predictions

Because the sequential gap filling approach produces different results depending on the order of gap filling, we chose to maintain many possible structures resulting from random permutations of the input media conditions rather than select a single GENRE structure for downstream analysis. Not knowing the “true” network structure, we considered each different structure to be a “hypothesis” and analyzed them collectively. For each of 2 to 30 training media conditions we produced 21 GENREs by randomizing the gap fill order (Supplemental Fig 1). We then evaluated each GENRE individually by predicting growth or no growth on 34 test media conditions (17 media conditions which experimentally supported growth and 17 which did not) using flux balance analysis (FBA). We found that each GENRE produced slightly different growth predictions, resulting in some GENREs being more accurate than others (Fig 4). In order to generate predictions using the ensemble, we treated each GENRE’s prediction as a single vote, and pooled the votes using a threshold (Fig 1B). We tested three qualitatively different thresholds; “any”, “majority”, and “consensus”. The “any” threshold simply requires that at least one GENRE predict growth in a particular media condition. The “majority” threshold requires greater than half to predict growth, and the “consensus” threshold requires all GENREs to predict growth. We evaluated the growth predictions in terms of accuracy, precision, and recall (see Materials and Methods).

**Fig 4.**
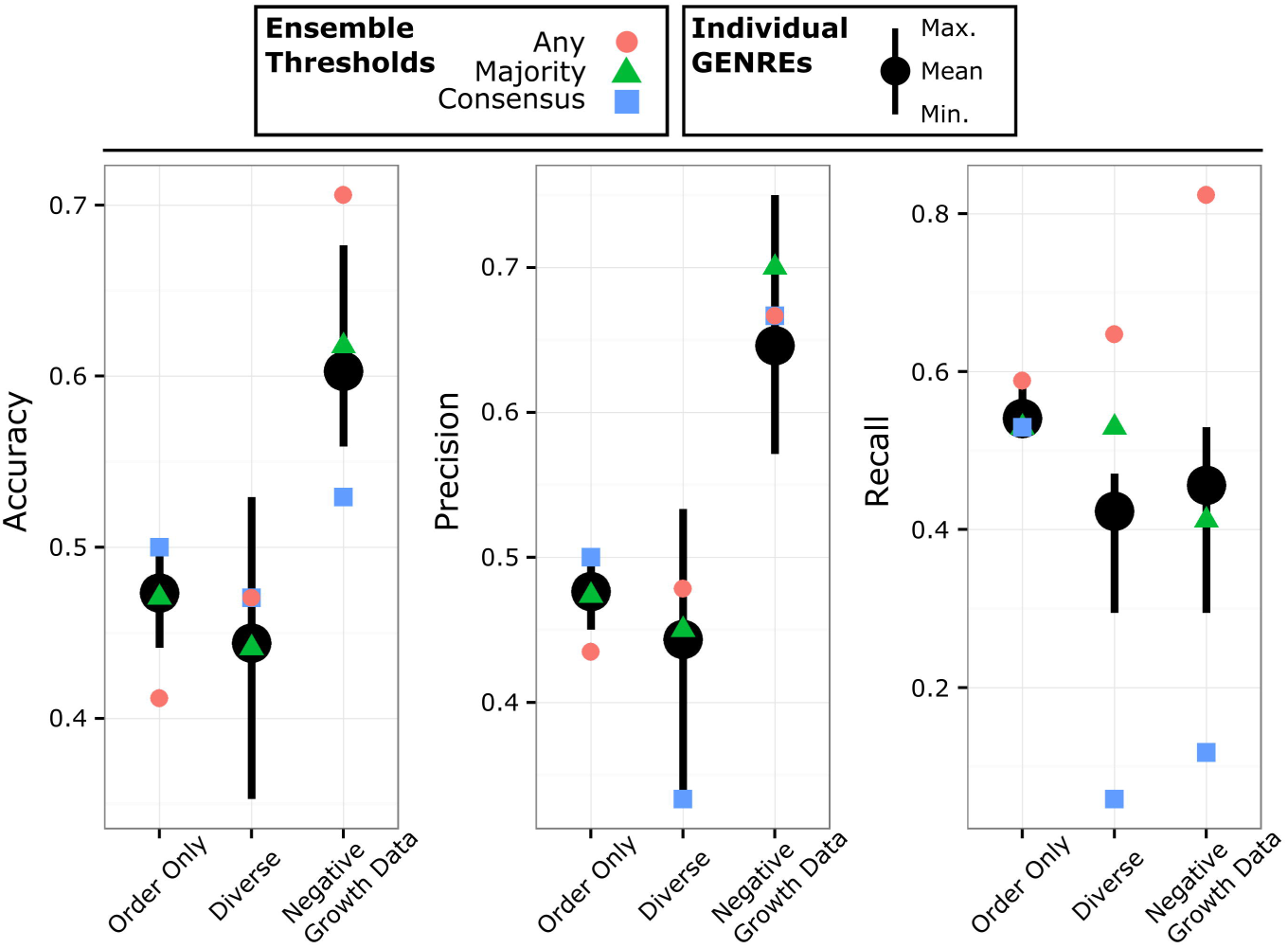
Ensembles generated by gap filling against the same media conditions in different orders. Using 20 media conditions, we generated 21 GENREs, where each GENRE was gap filled using either: a different order of the same input media conditions (“Order Only”), random weighting of reactions in database and random subsets of reactions from draft (“Diverse”), or a diverse ensemble which also included negative growth data through a trimming step (“Negative Growth Data”). We evaluated the accuracy, precision, and recall of every individual GENRE and of the ensembles by predicting growth on 17 positive media conditions and 17 negative media conditions which were not used during gap filling. The average of the individual GENREs is shown as black points with the maxima and minima as black lines extended above and below. The ensemble predictions using the three different thresholds are shown as red circles “any”, green triangles “majority”, and blue squares “consensus”. Note that there is less ensemble diversity when differences result only from media condition ordering (maxima/minima of “Order Only” compared to “Diverse” or “Negative Growth Data”). Adding additional diversity results in GENREs with both greater and lower accuracy than the best and worst of “Order Only”. Addition of the trimming step (“Negative Growth Data”) improves overall accuracy and precision by ~15%. In terms of ensemble thresholds, the “majority” threshold tends to perform similarly to the average of the individual GENREs. The “any” threshold achieves recall as good or better than the best individual GENREs. The “consensus” threshold performs consistently well in terms of accuracy and precision if there is very little diversity in the ensemble (“Order Only”).

We found that the “majority” threshold led the overall ensemble to achieve average accuracy with respect to the individual GENREs, consistently outperforming the least accurate of the individual GENREs (Fig 4 “Order Only”). The “any” threshold decreased overall accuracy and precision to be worse than any individual GENRE, but increased the recall to match the best individual GENREs. At the other extreme, we found that the “consensus” threshold led to accuracy and precision that matched the very best individual GENREs but diminished recall.

### Increasing the Diversity of Network Structures Increases Recall

While different gap fill order does result in different GENRE structures, the differences are relatively small (tens of differences relative to hundreds of reactions overall). In order to span a greater range of potential GENRE structures, we added random weights (drawn from a uniform distribution) to the reactions in the gap filling step (see Materials and Methods). The rationale is that given two pathways of slightly different length but the same biological function, random weights will occasionally favor the longer pathway, thus exploring alternatives that would otherwise be unobserved given a strictly parsimonious procedure. Additionally, each GENRE was reconstructed using a random subset of only 80% of the reactions from the draft GENRE from the Model SEED. Using this new procedure, we reconstructed ensembles of 21 GENREs using 2 through 30 training media conditions (Supplemental Fig 2). We evaluated the accuracy by predicting growth on the same 34 test media conditions as before. The resulting accuracy, precision and recall of the individual GENREs were essentially the same on average (Fig 4 “Diverse”), but the distribution spanned a much greater range, both positively and negatively. In this case, the “majority” threshold again achieved average behavior with respect to the individual GENREs, outperforming the least accurate individual GENREs (Fig 4 “Diverse”). The “any” threshold tended to achieve the best accuracy and precision, although not quite as good as the best individual GENREs. However, the “any” threshold achieved the best recall, better than the best individual GENREs and better than the recall achieved with a less diverse ensemble.

### Accounting for Negative Growth Conditions Greatly Improves Ensemble Accuracy, Precision and Recall

Our experimental growth data for *P. aeruginosa* UCBPP-PA14 included both positive (media conditions which supported growth) and negative results (media conditions which did not support growth). We formulated an optimization-based procedure which allowed us to incorporate information inherent in negative growth conditions into our automated curation (see Materials and Methods). In brief, the optimization problem identifies a minimal number of reactions to “trim” from a GENRE in order to prevent growth on negative media conditions while maintaining growth on positive media conditions. As before, we generated ensembles of 21 GENREs for 2 through 30 positive media conditions (Supplemental Fig 3). We used random reaction weights, random subsets of 80% of the reactions from the draft GENRE from Model SEED, and this time we selected 10 negative media conditions for each GENRE (distinct from the negative conditions used to assess accuracy) and incorporated them using our trimming procedure. We found that incorporation of the negative media conditions increased the accuracy and precision of both the individual GENREs and the ensembles by ~15% (Fig 4 “Negative Growth Data”). The “majority” threshold once again approximately tracked the average GENRE accuracy, precision and recall. The “any” threshold achieved accuracy and recall that were often better than the top individual GENREs, with recall exceeding that achieved previously.

### Ensembles Achieve Greater Precision or Recall Than Best Individual GENREs When Predicting Essential Genes

We evaluated the ability of ensembles to predict gene essentiality. We generated an ensemble of 51 GENREs, each created by gap filling with a random subset of 25 of the total 47 positive media conditions (53%), 10 randomly-selected negative media conditions from the total of 40 (25%), and 1,210 randomly-selected reactions from a total of 1,512 in the draft GENRE generated by Model SEED (80%). Genes were associated with reactions based on the assigned gene-protein-reaction (GPR) relationships from the draft GENRE. We used an *in silico* representation of CF sputum medium and predicted gene essentiality by removing reactions associated with each gene in turn (according to the GPR logic) and running FBA. We compared the resulting gene essentiality predictions with experimental results [19]. We found that the “majority” threshold resulted in better accuracy and recall than the average of individual GENREs, and drastic improvement over the worst GENREs (Fig 5). The “consensus” threshold resulted in a ~20% increase in precision over the best individual GENRE and greater than 100% increase over the worst individual GENRE. Unsurprisingly, the increased precision of the “consensus” threshold comes at the cost of reduced recall. The “any” threshold resulted in lower precision but a ~40% increase in recall over the best individual GENRE and a ~170% increase over the worst individual GENREs.

**Fig 5.**
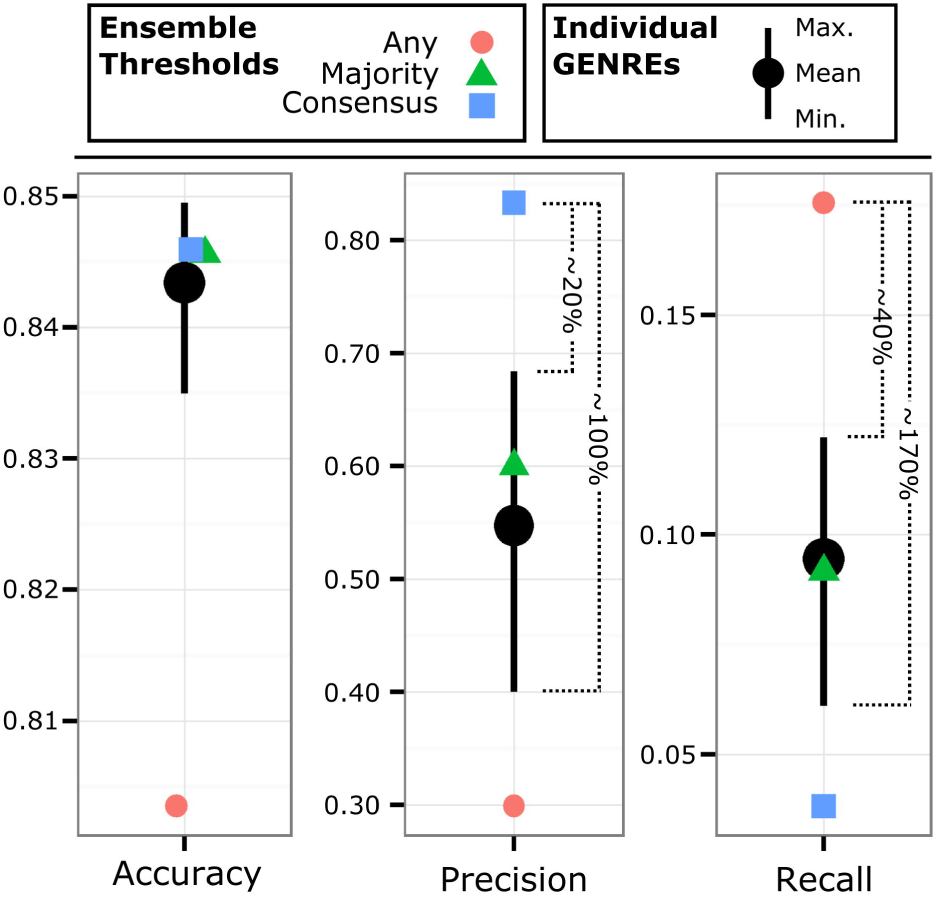
Ensembles outperform individual GENREs when predicting gene essentiality. We generated an ensemble of 51 GENREs by gap filling against 25 randomly-selected positive growth conditions, 10 negative growth conditions, and 80% of the reactions from the Model SEED draft network. We predicted gene essentiality in CF sputum medium and compared the predictions to *in vitro* gene essentiality data. We found that the “consensus” threshold (blue squares) achieved a ~20% increase in precision over the best individual GENRE and a ~100% increase in precision over the worst individual GENRE. Similarly, the “any” threshold (red circles) achieved a ~40% increase in recall over the best individual GENRE and a ~170% increase over the worst. Note the threshold-dependent tradeoff between precision and recall.

### Increasing Ensemble Size Improves Predictions for Small Ensembles

Using the same ensemble of 51 GENREs from above, we examined the effect of ensemble size on predicting essential genes. We sampled with replacement 10,000 small ensembles from among the 51 GENREs for ensemble sizes of 2 through 51. We evaluated the accuracy, precision and recall against the same gene essentiality data set using a “majority” threshold. We found that smaller ensembles were less accurate, less precise, and more variable than larger ensembles (Figs 6A and 6B). Increasing size improved predictions but with diminishing benefits as the ensemble grew larger. Interestingly, with this “majority” threshold, average recall increased initially, but diminishes again as the ensemble grows larger (Fig 6C).

**Fig 6.**
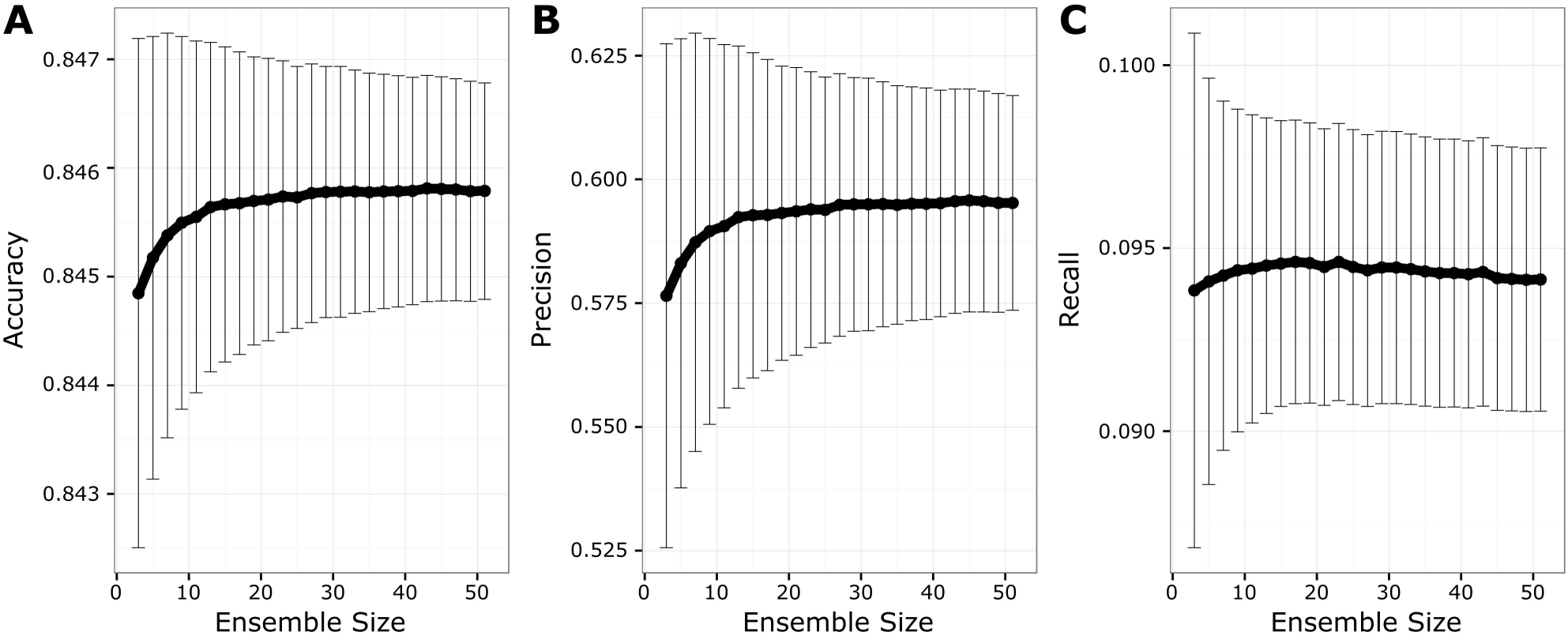
Increasing ensemble size improves performance initially. Using the same ensemble of 51 GENREs, we used bootstrap sampling to simulate 10,000 ensembles of sizes 2 through 51. We evaluated the performance of each sampled ensemble in terms of accuracy (**A**), precision (**B**), and recall (**C**) on the gene essentiality predictions using the “majority” threshold. We found that accuracy and precision increased sharply until around 15 GENREs, at which point gains were less pronounced with additional GENREs. This result suggests that increasing ensemble size does not infinitely improve ensemble performance.

### Common Reactions in Ensemble Are Consistent with Manually-Curated Reconstruction

In order to characterize the way gap filling distributes reactions throughout the ensemble, we generated an ensemble of 100 GENREs (Fig 7A). Each GENRE was reconstructed using a randomly-selected 80% of the reactions in iPAU1129, and then sequentially gap filled from the independent, universal reaction database using 25 random positive growth conditions. We found that before gap filling, the “correct” reactions from iPAU1129 were initially distributed in a bell curve throughout the ensemble (Fig 7B). The vast majority of “correct” reactions were found in 50 or more of the GENREs, and in 80 GENREs on average. In contrast, the “incorrect” reactions (those added by gap filling but which were not in the original iPAU1129) were distributed sporadically, with the majority being found in 10 or fewer GENREs. After the gap filling step, 65 “correct” reactions were found to have been added to every GENRE, suggesting a core set of “correct” reactions that were required for biomass production in any condition. We observed that the most common reactions (found in 50 or more GENREs) were overwhelmingly “correct” reactions from iPAU1129 (Fig 7C). All of these most common reactions (both “correct” and “incorrect”) were involved in the production of biomass components, particularly amino acids.

**Fig 7.**
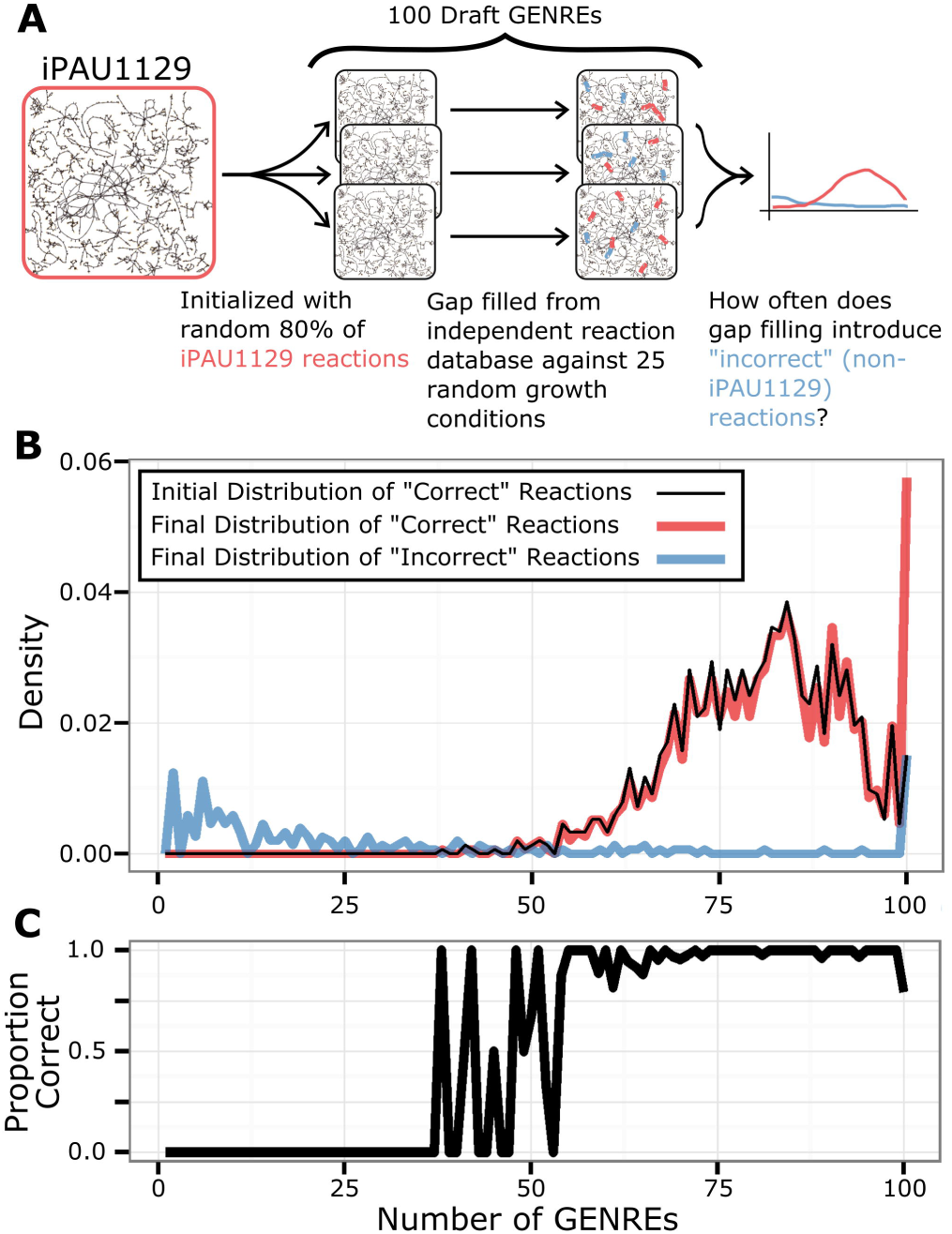
Common reactions in ensemble are consistent with manually-curated reconstruction. **A**. We generated an ensemble of 100 GENREs using a randomly-selected 80% of the reactions in iPAU1129, and then sequentially gap filled from the universal reaction database using 25 randomly-select positive growth conditions. **B**. The distribution of reactions throughout the ensemble is displayed as the proportion of reactions (y-axis) which are found in a given number of GENREs (x-axis). “Correct” reactions are whose which are found in the manually-curated iPAU1129, while “incorrect” reactions are those which are added during gap filling but not found in iPAU1129. We observed that there is a common set of reactions which were found in all 100 GENREs. The majority of this common set are “correct” (88 reactions) while 23 are “incorrect”. **C.** The common reactions (found in 50 or more GENREs) consist of a greater proportion of “correct” reactions. “Incorrect” reactions tend to be uncommon.

### Identifying Small Molecules Which Interact with Unique Streptococcus *Species*

We demonstrate how EnsembleFBA can be implemented in a systems biology workflow. We selected six species from the genus *Streptococcus* which all have growth phenotype data available through a previous study [20]. We reconstructed an ensemble for each species: *Streptococcus mitis*, *Streptococcus gallolyticus*, *Streptococcus oralis*, *Streptococcus equinus*, *Streptococcus pneumoniae* and *Streptococcus vestibularis* (Fig 8A). For each species, we generated a draft GENRE using the Model SEED online interface. We generated a diverse ensemble of 21 GENREs from each Model SEED draft, and gap filled each member GENRE using 25 random growth conditions specific to that species. We mapped all genes (translated to protein sequences) from each *Streptococcus* species to small molecule protein binding sequences from DrugBank using NCBI standalone BLASTP and an e-value threshold of 0.001 [21,22]. For all potential gene targets, we used the ensembles to predict gene essentiality using a “majority” threshold in rich media.

**Fig 8.**
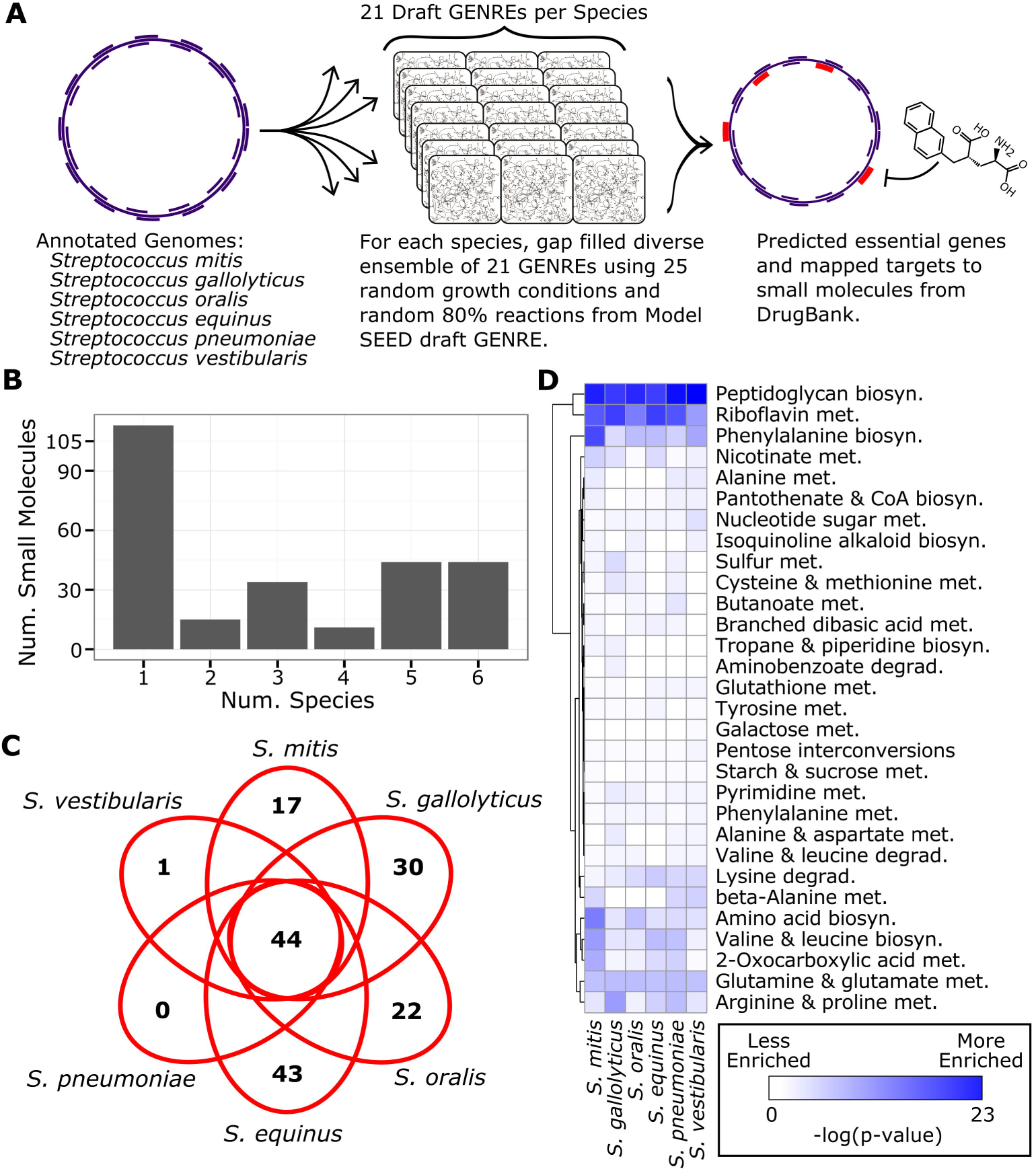
EnsembleFBA predicts unique essential genes targets of small molecules in six *Streptoccocus* species. **A**. We reconstructed ensembles of 21 GENREs for six *Streptococcus* species based on draft GENREs from the Model SEED and 25 growth conditions. From among all genes within all six species, we identified with potential binding interactions with small molecules from the DrugBank database, and used the ensembles to predict the essentiality of those genes. We found 261 small molecules with potential binding to essential gene products. **B**. Many small molecules interact with an essential gene in only one species, while a core set of 44 small molecules interact with essential genes in all six species. **C**. Distribution of small molecule interactions with essential genes, unique and conserved among the six species. Note that 44 small molecules interact with essential genes in all six species. *S. equinus* is predicted to have essential genes uniquely interact with 43 small molecules, while *S. pneumoniae* is predicted to not have any essential genes which interact with unique small molecules. **D**. Subsystem enrichment among essential reactions by species. We predicted reaction essentiality for all six species in rich media and then calculated a p-value indicating the likelihood of observing each subsystem among the essential reactions given the total number of reactions associated with that subsystem. For clarity we display the –log(p-value), where darker colors indicate greater enrichment (i.e. a disproportionate number of reactions in that subsystem are predicted to be essential). Note that some subsystems are enriched among essential reactions in all six species (e.g. Peptidoglycan biosynthesis) while others are uniquely enriched in a specific species (e.g. Phenylalanine biosynthesis in *S. mitis*).

We found 261 small molecules in DrugBank that potentially bind to the products of 169 essential genes (evenly distributed throughout the six species). Many of these small molecules (113) interact with an essential gene in only one of the species, while 44 were predicted to target conserved essential genes in all six species (Fig 8B). *S. equinus* was predicted to have the most essential genes interact with unique small molecules while *S. pneumoniae* was not predicted to have essential genes interact with any unique small molecules (Fig 8C). As an example of a conserved small molecule interaction, DB04083 (N’-Pyridoxyl-Lysine-5’-Monophosphate) is predicted to interact with essential aspartate aminotransferases in all six species. Alternatively, DB03222 (2’-Deoxyadenosine 5’-Triphosphate) is only predicted to interact with an essential ribonucleotide reductase in *S. gallolyticus*.

To better understand the differences between the metabolic networks which underpin these small molecule screen results, we predicted reaction essentiality in rich media for all six species using a “majority” threshold. We found that several metabolic subsystems were enriched among essential reactions beyond what would be expected from random chance (Fig 8D). Some subsystems were enriched in all six species, such as Peptidoglycan biosynthesis, indicating that these reactions related to cell wall biosynthesis are disproportionately essential in all six species. Other subsystems were enriched among essential reactions in a unique species. For example, *S. mitis* is predicted to have a greater proportion of essential reactions related to Amino acid metabolism than other species, perhaps indicating that *S. mitis* has less redundancy in those pathways than the other six species. Essential reactions related to Butanoate metabolism were most enriched in *S. pneumoniae*, while essential reactions in Lysine degradation were most enriched in *S. equinus*. Interestingly, reactions associated with core metabolic functions (e.g. Amino acid biosynthesis, Valine and leucine biosynthesis, Phenylalanine biosynthesis) were not equally enriched among essential reactions for all species.

## Discussion

Genome-scale metabolic network reconstructions (GENREs) have been used for decades to assemble information about an organism’s metabolism, to formally analyze that information, and in so doing, to make predictions about that organism’s behavior in unobserved or unobservable contexts. A major barrier preventing more widespread use of GENREs, particularly in non-model organisms, is the extensive time and effort required to produce a high-quality GENRE. Many automated approaches have been developed which reduce this time requirement (e.g. Model SEED, GLOBUS, CoReCo, RAVEN) [13–15]. We demonstrate that gap filling—although our results apply to many automated curation approaches—can lead to many potential GENRE structures depending on the ordering of the input data. Rather than arbitrarily selecting a single GENRE from among many possible networks (which are all reasonably consistent with the available data), we found that collecting many GENREs into an ensemble improved the predictions that could be made. We call this approach “EnsembleFBA” and emphasize that ensembles are a useful tool for dealing with uncertainty in network structure. We demonstrated how ensemble diversity impacts predictions. We show that EnsembleFBA correctly identifies many more essential genes in the model organism *P. aeruginosa* UCBPP-PA14 than the best individual GENREs. We showcase how EnsembleFBA can be utilized in a systems biology workflow by predicting how small molecules interact with different essential genes in six *Streptococcus* species. Ensembles increase the quality of predictions without incurring months of manual curation effort, thus making GENRE approaches more practical for applications which require predictions for many non-model organisms. We have provided code to facilitate the creation and analysis of ensembles of GENREs.

Gap filling is a common step during the GENRE curation process, both for manually- and automatically-curated GENREs [5]. We used a linear (rather than binary) gap filling algorithm to expand GENREs so that they are capable of producing biomass *in silico* on growth media which supports growth of the organism *in vitro*. Gap filling algorithms suggest parsimonious reaction sets from some “universal” biochemical database which, if added to a GENRE, will allow growth in the new environment [6,17,18,23]. Often, multiple reaction sets can enable growth, so some heuristics are needed to select a final solution. Sometimes gene homology metrics are used to select a solution, such that genes which catalyze the suggested reactions are compared to the current genome, and the reaction set with the best matches in the current genome are selected as the final solution. When validated, these solutions can lead to re-annotation of the genome [6]. During automated curation, there is less opportunity for extensive validation, and so the first or the most parsimonious solution is selected. As we demonstrated, the order of gap filling can change the final outcome, thus producing GENREs with different structures from the exact same input data (Fig 2). Under these circumstances, it is difficult to know which solution is most correct without additional data.

A possible way around this issue of gap fill order is to remove the sequential nature of gap filling entirely and use a global gap filling approach. We demonstrate that not only is such a global approach much slower (quadratic increases in solution time as growth media conditions are added), but the solutions are no more parsimonious or biologically relevant (Fig 3). Alternatively, we found that two additional innovations improved the predictions that could be obtained from automatically-generated GENREs: the addition of negative growth conditions and the collection of multiple GENREs into ensembles.

Negative growth conditions have not been extensively incorporated into GENRE curation. To our knowledge, only one group has developed an approach for removing reactions in order to prevent growth in specific conditions [23]. In that case, the reactions were not removed from the GENRE, but rather, prevented from carrying flux under particular conditions. This approach was supported by a biological justification that certain enzymes may not be functional under certain conditions [23]. Our approach is different in that it seeks to produce a single GENRE structure that is consistent with all available data, positive and negative. We achieved this by removing a minimal reaction set to simultaneously prevent growth in negative conditions and allow growth in positive growth conditions (see Materials and Methods). By utilizing this untapped source of information, we found that average GENRE accuracy increased by ~15% (Fig 4). Automatically incorporating negative growth conditions is a little-explored area that has the potential to make better use of growth screening data.

Ensembles have been used for many years in the machine learning community to leverage the strengths of many different models to improve predictions [24,25]. Ensembles have been used previously to analyze GENREs from a kinetic standpoint [26]. Because kinetic parameters are usually unknown for an entire genome-scale network, ensembles of kinetic parameters are generated such that all parameter sets lead to the same steady state [26]. In this way, ensembles can represent the space of allowable kinetic parameters. Our approach to generating ensembles is different in that we attempt to represent the space of allowable GENRE structures rather than kinetic parameters.

Ensembles provide a significant advantage over individual GENREs by tuning for specific results with defined decision thresholds (Figs 4 and 5). Consistently, by using the “any” threshold, recall can be made to equal or exceed the best individual GENREs. This result makes sense, considering that different network structures will result in different growth or gene essentiality predictions. By accepting any essential gene prediction from among the constituent GENREs, we cast a wider net and capture many more of the true essential genes and growth conditions. The fact that many individual GENREs contribute unique but true predictions suggests that each GENRE recapitulates elements of the “true” network structure (Fig 7). Similarly, by using the “majority” threshold, the ensemble predictions perform like the average GENRE (Figs 4 and 5). By requiring a majority of GENREs to agree, the ensemble guards against poor predictions and, in most cases, outperforms the worst individual GENREs. Finally, if precision is the overall goal, a “consensus” threshold provides confidence that the majority of positive predictions are true positives (Fig 5).

We observed that ensemble performance is limited by the quality of the GENREs which form the ensemble. The choice of decision threshold (“any”, “majority”, or” consensus”) did not consistently improve overall accuracy of the ensemble. However, by improving the individual GENREs using negative information, the overall ensemble accuracy improved dramatically (Fig 4). Also, it should be noted that the computational burden required by ensembles will always be greater than the burden of a single GENRE. For all the examples in this study, computational burden scales linearly with the number of GENREs in the ensemble (ensemble of size N GENREs will require N times longer to calculate FBA solutions) which is a modest expectation in practice. Other applications, like predicting species interactions, would not scale linearly if all possible pairs of GENREs between two ensembles were simulated.

Increasing ensemble diversity impacted ensemble recall, but did not have an obvious effect on overall accuracy. Some degree of diversity is required in order to gain any advantage through an ensemble representation. In the “Order Only” ensemble (generated simply by changing the order of gap filling; Fig 4) there were only small differences between any of the GENREs so it was difficult to improve on the best GENRE. By injecting greater diversity through random weights and random subsets of the data, we observed much greater variation in individual GENRE performance (both positively and negatively), but the average accuracy was the same as the low diversity ensemble (Fig 4). The advantage of diversity is in casting a wide net and thus improving ensemble recall, particularly when combined with an “any” decision threshold. In practice, the choice to increase diversity or not will depend on the goals of the analysis. If the goal is to generate many candidate essential genes or media conditions, then more diversity will be advantageous. If the goal is to generate fewer, more confident predictions, then minimizing diversity will be most effective.

EnsembleFBA is easily integrated into systems biology workflows. As an example, a current challenge in systems biology is to identify species-specific drug targets so that therapies will not disrupt the healthy microbiome structure [27,28]. We reconstructed ensembles for six *Streptococcus* species by gap filling with growth phenotype data, we predicted essential genes and mapped those genes to potential small molecule binding partners within a matter of hours, and can have more confidence in the quality of the gene essentiality predictions than if we were to work with single GENREs for each species (Fig 8). The process scales well with the number of species, such that 12 or 100 species would not take significantly longer than six, and the quality of the predictions is maintained with scale. It is interesting to note that among *Streptococcus* species, there are generally small molecules which can be selected to uniquely interact with essential genes in a single species, and other small molecules which interact with conserved essential genes (Fig 8C). The observed interactions between essential genes and small molecule ligands are species-specific because of differences in network structure which lead to some metabolic subsystems being disproportionately represented among essential reactions (Fig 8D). In the search for species-specific drug targets, it is important to consider, not only the presence or absence of a particular gene, but also the role of that gene in the broader network context, and improved systems biology tools such as EnsembleFBA can help to elucidate that context with greater confidence.

Gap filling is not the only GENRE reconstruction approach that produces many possible solutions. Likelihood-based gap filling produces a distribution of possible annotations for each gene in a genome, assigning a probability to each [14,16,29]. Network structure is then based on maximizing the likelihood over all possible solutions. Ensembles could be generated easily using this type of framework by sampling many alternative solutions around the maximum likelihood. Indeed, it may be beneficial to create an ensemble using GENREs reconstructed using several different methods. We suggest that there are many possible ways to generate ensembles such that they will allow researchers to generate better predictions about under-studied organisms.

Finally, we foresee ensembles playing an important role beyond improving predictions, for example, in experimental design and model reconciliation. Within a diverse ensemble, many possible network structures are represented, and it is expected that some structures will be closer to the truth than others. We suggest that ensembles can be leveraged to design an optimal series of experiments to weed out the most incorrect network structures. For instance, such an approach could select the most differentiating carbon sources to experimentally test, or the most differentiating essential genes. Ensemble-guided experimental design could save time and experimental resources. Model reconciliation is another field that could benefit from ensembles [8,30]. Given GENREs for two different species, reconciliation is the process of removing systematic differences from the two GENREs so that any differences which remain are due to biology alone. Systematic differences often result from arbitrary choices during the process of reconstruction. Ensembles could be used to automate the reconciliation process by representing the space of possible GENREs for each species and the reconciled versions would be the two models from the two spaces that are most similar to each other. Thus, ensembles have potential to improve other tasks than prediction, including experimental design and mapping the space of GENRE structures for tasks like reconciliation.

## Materials and Methods

### Code and Data Availability

All data, Matlab (Natick, MA, USA) implementations of algorithms, Matlab simulation scripts, results files and figure generation scripts are publically available in our online repository: https://github.com/mbi2gs/ensembleFBA

### Data Sources

All biochemical reference data was obtained from the Model SEED database (https://github.com/ModelSEED/ModelSEEDDatabase). The metabolic reaction and compound databases were parsed and formatted for use in Matlab using a custom Python script available in our repository (“format_SEED_data.py”). A draft network for *P. aeruginosa* UCBPP-PA14 was automatically generated using the Model SEED web service (http://modelseed.org/genomes/). Similarly, draft networks were generated for *Streptococcus mitis* ATCC 6249, *Streptococcus gallolyticus* ICDDRB-NRC-S3, *Streptococcus oralis* ATCC 49296, *Streptococcus equinus* AG46, *Streptococcus pneumoniae* (PATRIC ID 1313.5731), and *Streptococcus vestibularis* 22-06 S6.

Representations of media conditions (including minimal media and cystic fibrosis sputum medium), and biomass representations were drawn from previous GENRE analyses of *Pseudomonas aeruginosa* [31,32].

*P. aeruginosa* PA14 essential genes in cystic fibrosis sputum medium were experimentally identified previously [19].

A manually curated, and thoroughly validated GENRE of *P. aeruginosa* UCBPP-PA14 called iPAU1129 was developed previously (Bartell et al. In review), along with Biolog growth screen data for *P. aeruginosa* UCBPP-PA14 indicating many media conditions in which this strain will and will not grow.

Growth phenotype data for six *Streptococcus* species was obtained from the file “Supplementary Data 1” of [20]. Small molecule amino acid binding target sequences were downloaded from the DrugBank website (http://www.drugbank.ca/) [21]. After identifying homologous genes to the target sequences using BLASTP [22], we used a custom python script to parse the results for input into Matlab (“listPossibleTargets.py”, available in repository).

### Linear Gap Filling

We implemented a linear (as opposed to binary) gap filling algorithm in Matlab, based on the algorithms FASTGAPFILL and FastGapFilling [17,18]. We used the Gurobi solver version 6.0.5 for all optimization tasks (Gurobi, Houston, TX, USA). To begin, we provide the algorithm with a universal database of metabolic reactions U, a universal database of exchange reactions X, a biomass reaction, and a set of growth conditions formatted as lower bounds on exchange reactions. The algorithm identifies a set of reactions from U and X that allow flux through the biomass reaction under all growth conditions. The algorithm is implemented as a linear program (LP) that minimizes the sum of the absolute value of all fluxes through U and X. The optimization problem takes the form:

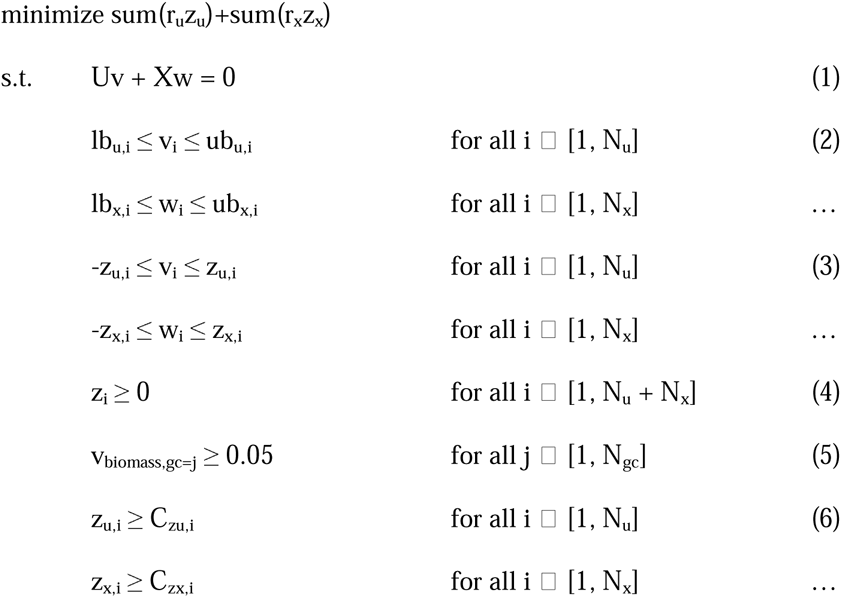

Where:

U is the universal reaction library (as a stoichiometric matrix)
X is the universal exchange library (same metabolites as U)
N_u_ and N_x_ are the number of reactions in U and X, respectively
N_gc_ is the number of growth conditions
v is the vector of fluxes through U
w is the vector of fluxes through X
lb_u,i_, ub_u,i_, lb_x,i_, and ub_x,i_ are the lower and upper bounds on v_i_ and w_i,_ respectively v_biomass,gc=j_ is the flux through the biomass reaction under growth condition j C_zu,i_ and Cz_x,i_ are variables that can force reactions from U and X to be included z is a continuous variable which acts as a constraint on the absolute values of elements in v and w.
r is an optional weight on z which can be randomized

In order to incorporate genome annotations from a specific organism, we force the inclusion of all associated reactions from those annotations using the C_zu_ variables. Note that unlike a binary optimization, the LP minimizing the sum of the absolute flux values through U and X does not necessarily result in a solution with the fewest reactions, but rather the solution which requires the minimum sum of the absolute values of the fluxes through it. The LP here can be extended to utilize multiple growth conditions simultaneously (global approach) by duplicating the U and X matrices, once for each growth condition, but minimizing a single set of z variables across all conditions. To gap fill using multiple growth conditions sequentially, we gap fill using the first growth condition, incorporate the solution into the GENRE, then repeat the process for all growth conditions. Our Matlab function “expand()” implements this optimization problem.

### Incorporating Negative Growth Conditions by Trimming Reactions

We implemented a binary optimization problem to trim minimal reactions from a GENRE in order to prevent growth under negative growth conditions while simultaneously maintaining growth in the positive growth conditions. As input to the algorithm, we provide a GENRE, and a set of both positive and negative growth conditions. We chose to run FBA first to identify mismatches between the computational predictions and the *in silico* data (negative growth conditions that erroneously predicted to support growth *in silico)*. Having identified those, we then ran FBA on all the positive growth conditions to identify the top five with flux distributions most similar to the flux distribution of the negative growth condition. The GENRE and the selected growth conditions are passed to the trimming problem, which takes the form:

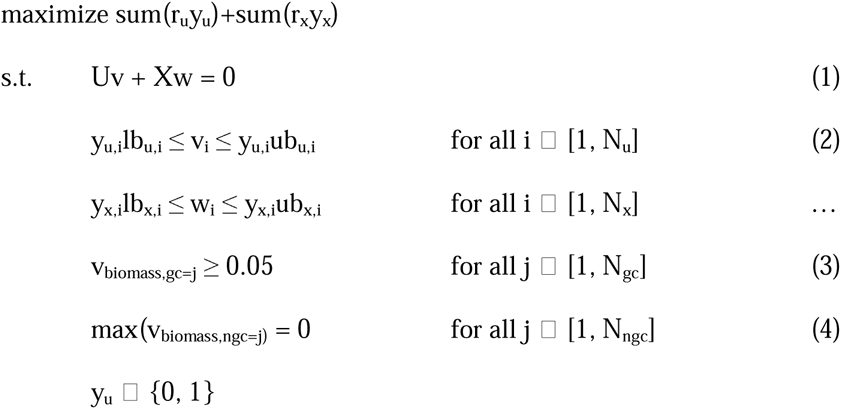

Where:

U is the set of reactions from the GENRE (as a stoichiometric matrix)
X is the set of exchange reactions from the GENRE (same metabolites as U)
N_u_ and N_x_ are the number of reactions in U and X, respectively
v is the vector of fluxes through U
w is the vector of fluxes through X
lb_u,i_, ub_u,i_, lb_x,i_, and ub_x,i_ are the lower and upper bounds on v_i_ and w_i,_ respectively v_biomass,gc=j_ is the flux through the biomass reaction under growth condition j y is a set of binary variables which determine inclusion of reactions in the network r is an optional weight on y which can be randomized

The term max(v_biomass,ngc=j)_ = 0 requires that the maximum possible flux through the biomass reaction for the negative growth conditions is constrained to be zero. In order to implement this constraint, we took advantage of duality theory as has been done previously [7]. Specifically, the optimal objective value of the dual of a linear program will equal the optimal value of the primal. By constraining the primal and dual objectives to equal each other, we can ensure that the flux through the biomass objective is maximized. We can replace the term max(v_biomass,ngc=j)_ = 0 with the following constraints:

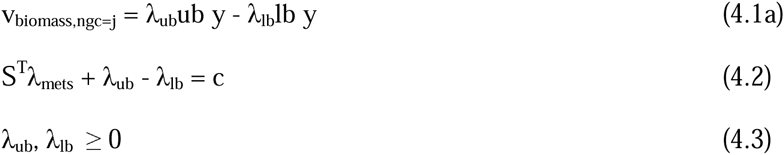

Where λ_mets_, λ_ub_ and λ_lb_ are the dual vectors associated with the metabolites, upper and lower bounds of the primal problem. Note that the terms λ_ub_ub y and λ_lb_lb y are quadratic, requiring a multiplication of the binary inclusion variable y with the dual variables. Because y is a binary variable, in this case the quadratic constraints can be converted to linear constraints through the use of additional variables:

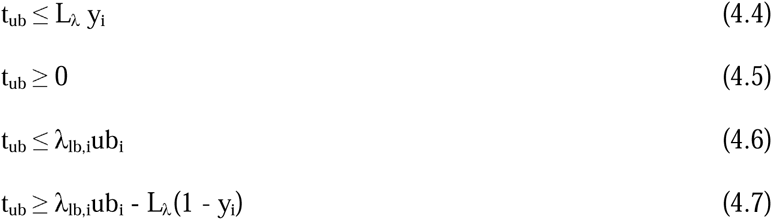

Where t_ub_ is a stand-in for the product λ_ub_ub y and L is a large number greater than or equal to the upper bound on λ_ub_ub y (e.g. 1000). Similar constraints would be produced for the product λ_lb_lb. The quadratic constraint above can then be replaced by a linear constraint:

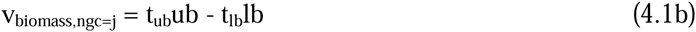

Our Matlab function “trim_active()” implements this optimization problem.

### Iterative Approach to Reconstructing GENREs Consistent with All Growth Screening Data

We implemented an iterative algorithm to integrate the LP expansion step with the binary trimming step. The algorithm first applies the expansion step to produce a GENRE that is capable of growing in all positive growth conditions. Next, the algorithm checks for negative growth conditions that allow for biomass flux and for any that do, applies the trim step as described above. The algorithm iterates between the expand and trim steps until either a completely consistent GENRE structure is identified, or it reaches a maximum attempts limit. A single attempt is completed if the GENRE structure is not yet consistent with the input growth conditions but stops making progress (stuck in a local optimum). In this case, a random reaction is removed from the GENRE and the search is re-initiated. If the maximum attempts limit is reached, the algorithm removes any negative growth conditions that are inconsistent with the positive growth conditions, and returns the final GENRE. This iterative algorithm is implemented in our Matlab function “build_network()”.

### Predicting Growth and Essential Genes

Growth media were simulated by setting the lower bounds on exchange reactions for the appropriate nutrients to negative values. The uptake of carbon source(s) limited the final flux through biomass. “Growth” was determined by maximizing flux through the biomass objective. We predicted “growth” if a positive, non-zero flux could be achieved through biomass. Gene knock-outs were simulated by generating a new GENRE which was missing the reactions dependent on the knocked-out gene. The reaction-gene dependence was determined by evaluating the binary logic of the GPRs provided by Model SEED. Our custom script to evaluate GPR logic is “simulateGeneDeletion()”.

We evaluated the growth predictions in terms of accuracy (TP + TN) / (TP + FP + TN + FN), precision (TP / TP + FP), and recall (TP / TP + FN) where TP = number of true positives, FP = the number of false positives, TN = the number of true negatives and FN = the number of false negatives. Precision indicates the number of positive predictions which are true positives. Recall indicates the number of positive events which were correctly predicted by the method.

### Predicting Small Molecule Interactions

We downloaded the Drug Target Sequences for small molecules in FASTA format from DrugBank [21]. Using NCBI standalone BLASTP and an e-value cutoff of 0.001, we identified homologous sequences in all six *Streptococcus* proteomes [22].

### Metabolic Subsystem Enrichment

We downloaded KEGG subsystem annotations for the reactions in the Model SEED database (“KEGG.pathways.tsv”). After predicting essential reactions for each *Streptococcus* species, we used the hypergeomtric distribution to calculate the probability of drawing k essential reactions and finding that x or more are annotated with subsystem *j,* from a population of size M reactions, of which N are annotated with subsystem *j*.

### Computational Resources

The majority of our reconstructions and simulations were performed on a 64-bit Dell Precision T3600 Desktop computer with 32 GB RAM and eight 3.6 GHz Intel Xeon CPUs, running Windows 7. Incorporating negative growth information often lead to longer reconstruction times (sometimes 2 hours per GENRE) due to the binary optimization step. To accelerate the reconstruction time while incorporating negative growth information, we used the University of Virginia’s High Performance Computing Cluster.

### Generating Ensembles and Making Predictions Using Our Software

Our Matlab scripts for generating an ensemble (using the gap filling approach described in this work) and for analyzing an ensemble are freely available in a github repository (see Code and Data Availability). The Gurobi solver is required, in addition to our Matlab scripts. We have also included a tutorial script to guide the user through the necessary steps to generate and analyze an ensemble (“test_eFBA.mat”).

## Acknowledgements

The authors would like to thank Jennifer Bartell for providing a copy of iPAU1129 and the accompanying carbon source utilization data.

